# Mapping tumor microenvironment heterogeneity of human melanoma metastases across distant organs

**DOI:** 10.64898/2026.08.11.743891

**Authors:** Carlos Ariel Pulido-Vicuna, Marek Piprek, Philip Georg Demaerel, Oliver Bechter, Peter Vermeulen, Francesca Maria Bosisio, Jean-Christophe Marine, Joanna Pozniak

## Abstract

Distant metastasis has a major impact on melanoma mortality, yet it remains poorly defined how disseminated cells adapt to different host organs and their microenvironment. Here we applied single-nucleus RNA sequencing (snPATHO-seq) to archival FFPE melanoma metastases spanning brain, liver and lung from different patients. We built an atlas resolving malignant, immune and stromal compartments in each organ. Within the malignant compartment, we observed a differentiation axis of melanocytic, transitory, neural crest-like and neural crest-like/mesenchymal states; and meta-programs: interferon-responsive, hypoxic, cycling and stress. The immune compartment showed organ specificity: resident macrophage identity recapitulated the host tissue (microglia, Kupffer cells and alveolar macrophages). Brain-infiltrating myeloid and lymphoid cells were transcriptionally the most immunosuppressive and most cytotoxic, respectively, of the three sites. Non-malignant stromal populations, including fibroblasts, endothelial cells and pericytes, also carried distinct organ-specific transcriptional programs, suggesting that organ-of-residence effects extend beyond the malignant cells to the whole metastatic ecosystem. We provide a first multi-organ atlas of melanoma distant metastasis allowing deeper understanding of how a melanoma is remodelled by, and/or remodels, three distinct human organ environments.

## Introduction

Cutaneous melanoma mortality is largely driven by its high metastatic potential and dissemination to vital organs. Among these sites, the liver, lung and brain are the most frequently colonized [1]. Patients with distant metastases benefit the least from current standard of care therapies [2], which include targeted therapy and immune checkpoint blockade (ICB). Understanding the biology of established visceral and intracranial metastasis is crucial to improve outcomes for these patients.

Cancer evolution and therapy resistance in melanoma are partially driven by intratumour heterogeneity and phenotypic plasticity [3]. Single-cell profiling has shown that melanoma cells move along a plastic transcriptomic continuum of lineage-specific states that spans melanocytic, neural crest-like and neural crest-like/mesenchymal-like phenotypes [4–6]. Melanoma cells can also activate extrinsic transcriptomic metaprograms in response to microenvironmental cues such as immune infiltration or hypoxia [6,7]. This plasticity contributes to metastatic competence and adaptive ICB resistance [3,8].

This transcriptomic framework has been assembled almost entirely from primary cutaneous tumours, skin lesions and lymph-node metastases [6,9]. The most common and lethal visceral sites, particularly the liver and lung, remain largely unexplored at single-cell resolution [10]. Few studies have profiled brain metastases, but it is still unclear whether host organ-specific melanoma cell states exist [10,11]. Resolving whether the host organ imposes distinct melanoma cell states and meta-programs is central to understanding metastatic establishment, maintenance and therapy resistance.

There is a technical reason why the study of visceral metastases has been challenging compared to primary tumours. Conventional single-cell RNA sequencing requires prospectively collected, viable fresh tissue, which is rarely available for visceral metastases that are rarely resected [12]. To address this, we applied the snPATHO-seq protocol, which extracts single nuclei directly from archival formalin-fixed paraffin-embedded (FFPE) blocks [12]. This allowed robust single-nucleus RNA sequencing (snRNA-seq) of clinically annotated visceral and intracranial metastases.

Here we present a single-cell-resolution characterisation of human melanoma metastases from the brain, lung and liver. Profiling 22 patient samples, we analysed 132,054 nuclei, of which 122,555 were confidently assigned to the malignant, immune or stromal compartments. Bioinformatic analyses revealed both shared and organ-specific melanoma cell states and meta-programs across host organs. This atlas serves as a valuable resource for the melanoma community and beyond.

## Results

To characterise the cellular composition of distant melanoma metastases, we performed single-nuclei PATHO-seq (snPATHO-seq) [12] on formalin-fixed paraffin-embedded (FFPE) brain (n=5), liver (n=6) and lung (n=11) metastatic lesions from 20 different patients (Fig. 1A). Samples comprised treatment-naive biopsies (BT, n=12) and on-or post-treatment resections (OT, n=8); treatment status was unavailable for two liver samples (NA, n=2). Details about on-/post treatment samples derived from patients are detailed in Table S1.

**Figure 1.**
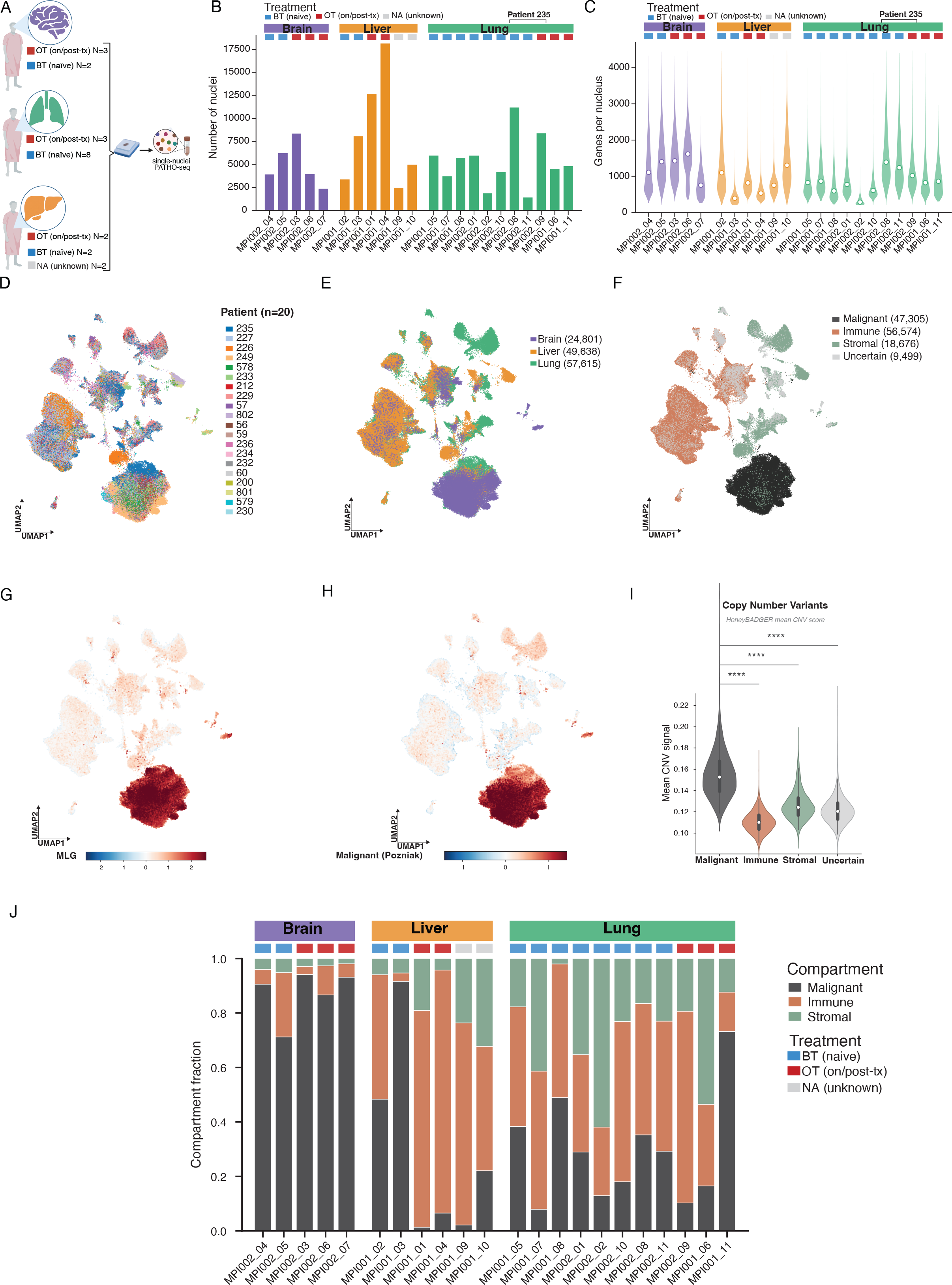
A cohort of distant melanoma metastases. **(A)** Study overview. Archival formalin-fixed paraffin-embedded (FFPE) metastatic melanoma samples from three organ sites: brain, liver and lung were profiled by single-nucleus PATHO-seq (snPATHO-seq). Per organ, samples are split by treatment status into treatment-naïve biopsies (BT) and on-/post-treatment resections (OT): brain (OT n = 3, BT n = 2), liver (OT n = 2, BT n = 2, NA n = 2) and lung (OT n = 3, BT n = 8). In total, 22 samples from 20 patients were analysed. **(B)** Number of nuclei recovered per sample after quality control, coloured by organ (brain, purple; liver, orange; lung, green) and ordered within each organ block. The coloured tiles above each bar indicate treatment status (BT, blue; OT, red; NA/unknown, grey). Samples from patient 235 are indicated. Recovery ranged from ∼1,400 to ∼18,100 nuclei per sample, with the lowest yields in brain. **(C)** Violin plots of the number of genes detected per nucleus for each sample, grouped and coloured by organ as in (B); white points mark the per-sample median. Gene-detection rates were comparable across organs despite differences in nucleus yield. **(D–F)** Uniform Manifold Approximation and Projection (UMAP) embeddings of all 132,054 nuclei computed on the scVI-integrated latent space (patient identity used as the batch covariate). Nuclei are coloured by **(D)** patient of origin (n = 20), **(E)** organ of origin (brain, 24,801; liver, 49,638; lung, 57,615 nuclei) and **(F)** assigned compartment: malignant (47,305), immune (56,574), stromal (18,676) and uncertain (9,499). UMAPs are shown as low-dimensional visual summaries only; compartment assignment was performed on the integrated latent space rather than on the two-dimensional embedding. **(G)** UMAP as in (D–F) coloured by the melanocytic minimal-lineage gene (MLG) signature score (derived from *PMEL, TYR, DCT, MLANA, MLPH, TFAP2A* and *PRAME*). High MLG signal is confined almost exclusively to the malignant compartment. **(H)** UMAP coloured by the malignant melanoma signature score (Poźniak et al.,Cell, 2024), which co-localises with the MLG-high region in (G). **(I)** Violin plots of the HoneyBADGER-derived mean copy-number-variant (CNV) signal per compartment. Malignant nuclei show significantly higher mean CNV signal than immune, stromal and uncertain nuclei. White points and boxes denote the median and interquartile range. **\*\*\*\***, p < 0.0001. **(J)** Stacked bar plots of the fraction of malignant, immune and stromal nuclei per sample, grouped by organ; treatment-status tiles (BT, blue; OT, red; NA, grey) are shown above each bar. Brain metastases carried a high malignant burden and a sparse tumour microenvironment (TME), whereas liver and lung metastases showed a more heterogeneous TME composition. Uncertain nuclei are excluded from this panel.

The number of nuclei recovered per sample varied from ∼1,400 to ∼18,100 (Fig. 1B). Between organs, the lowest number of nuclei was observed from the brain compared to liver and lung. In contrast, gene-detection rates were consistent across organs (Fig. 1C). The processing of the single-nuclei data involved several steps: per-nucleus quality control reduced the initial 163,616 nuclei to 154,102. We removed nuclei identified as doublets by a consensus of Scrublet [13], scDblFinder [14] and Solo [15], retaining 138,443 nuclei. At the pan-organ level, we integrated samples using scVI [16], considering the high inter-patient variability of this cohort [17,18]. While the provided UMAP plots serve strictly as low-dimensional visual summaries (Fig. 1D–F), following steps in the processing pipeline robustly resolved both common and tissue-specific cellular populations across all three metastatic sites. Among these, malignant cells were identified with a probabilistic classifier that combined melanoma, immune and stromal signature scores [6] including one derived from HoneyBADGER copy number variants (CNV) inference [19]. Malignant nuclei, as expected, showed higher mean CNV and melanoma signatures than immune and stromal populations (Fig. 1G,H,I). After removing ambiguous and ambient-contaminated nuclei, the final atlas comprised 132,054 nuclei divided into malignant (n=47,305), immune (n=56,574), stromal (n=18,676) and uncertain (n=9,499) compartments (Fig. 1F). Nuclei whose compartment posterior was ambiguous were retained as a distinct uncertain category; they are displayed in Fig. 1 but excluded from all downstream analyses, which use the 122,555 confidently assigned nuclei. Whereas brain samples showed a high malignant burden, and low TME, liver and lung metastases were marked by a more heterogeneous TME composition (Fig. 1J).

### Malignant cell characterisation across organ sites

To dissect the malignant compartment only, we used Harmony [20] because this tool was shown to be more suitable than scVI for the integration of less complex datasets [17]. We explored the transcriptomic heterogeneity on two levels (Fig. 2A), separating melanoma’s lineage-specific differentiation continuum from environmentally driven meta-programs and mitotic signal. This division was inspired by previous observations from our group and others [4–6, 9, 17, 21].

**Figure 2.**
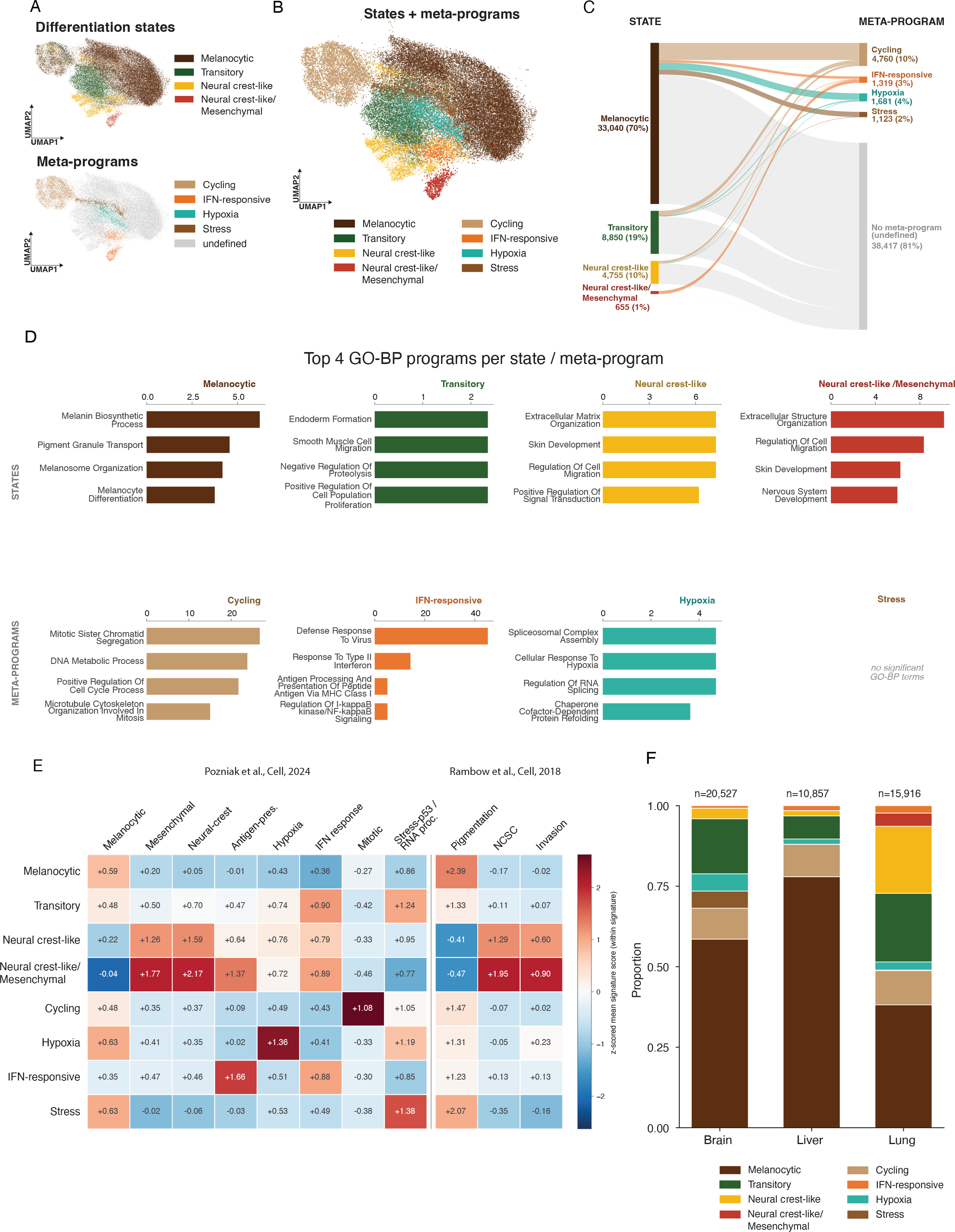
Malignant cell characterisation across organ sites. Analyses in this figure use the malignant nuclei (Harmony-integrated), with differentiation states assigned to 47,300 nuclei. **(A)** UMAP embeddings of malignant nuclei coloured, top, by lineage-specific differentiation state (melanocytic, transitory, neural crest-like, neural crest-like/mesenchymal) and, bottom, by environmentally driven meta-program (cycling, IFN-responsive, hypoxia, stress); nuclei not expressing a dominant meta-program are shown as undefined. **(B)** UMAP of malignant nuclei showing the combined assignment of differentiation states and overlaid meta-programs. **(C)** Sankey diagram relating differentiation state (left) to meta-program (right). State totals: melanocytic 33,040 (70%), transitory 8,850 (19%), neural crest-like 4,755 (10%) and neural crest-like/mesenchymal 655 (1%). Meta-program totals: cycling 4,760 (10%), IFN-responsive 1,319 (3%), hypoxia 1,681 (4%), stress 1,123 (2%); the majority of nuclei (38,417; 81%) carried no dominant meta-program. Cycling, hypoxia and stress activity was concentrated in melanocytic cells. **(D)** Top four enriched Gene Ontology Biological Process (GO-BP) terms for each differentiation state (top row) and each meta-program (bottom row), ranked by enrichment. Enrichment was computed on differential-expression markers after removal of organ-parenchyma ambient genes. The stress meta-program returned no significantly enriched GO-BP terms. **(E)** Heatmap of z-scored mean signature scores (scaled within each signature) for each differentiation state and meta-program (rows) across published reference signatures (columns; melanocytic, mesenchymal, neural-crest, antigen-presenting, hypoxia, IFN response, mitotic, stress-p53/RNA processing, pigmentation, NCSC and invasion; Poźniak et al., 2024 and Rambow et al., 2018). Concordance between reference signatures and the assigned annotations supports the malignant-cell classification. **(F)** Stacked bar plots of the proportion of malignant states and meta-programs by organ (brain, n = 20,527; liver, n = 10,857; lung, n = 15,916 nuclei). Brain and liver metastases were predominantly melanocytic, whereas lung metastases carried the largest dedifferentiated (neural crest-like and neural crest-like/mesenchymal) fractions.

We identified four lineage-specific melanoma cell states (melanocytic, transitory, neural crest-like, and neural crest-like/mesenchymal) and four metaprograms (cycling, hypoxia, stress, and interferon-responsive). For each cell, we assigned a differentiation state and determined whether specific metaprogram(s) were engaged (Fig. 2A, B). Note that most cells did not appear to express a dominant meta-program and were labelled as undefined (Fig. 2A bottom; Fig. 2C). By visualising the relationship between differentiation state and meta-programs we observed that the meta-program activity was unevenly distributed across states: the cycling, hypoxia and stress programs were expressed mainly by melanocytic cells (Fig. 2C). Functional enrichment of pathways supported the assigned identities as we observed known terms connected to each state and meta-program except for the stress population, which did not show significantly enriched terms. Consistent with previous observations, the melanocytic state was enriched in terms related to melanin biosynthesis and pigmentation. The transitory state showed both proliferation and cellular migration signals, neural crest-like state preferentially upregulated programs linked to extracellular matrix remodelling and skin development. Cells in the neural crest-like/mesenchymal state expressed similar programs to neural crest-like, nevertheless the extracellular matrix organization was enriched at a higher level (Fig. 2D) accompanied by the lowest expression of melanocytic gene signature, could imply a transition from neural crest, into an even more de-differentiated mesenchymal-like state. The four meta-programs recovered their expected biology. Enrichment was computed on differential-expression markers from which organ-parenchyma ambient genes had been removed, ensuring the recovered programs reflect malignant-intrinsic biology rather than tissue-of-origin contamination. To contextualize our findings, we used reference signatures from Pozniak et al. (2024) and Rambow et al. (2018) [5,6]. The strong correspondence between these established signatures and our defined annotations and gene ontology substantiates our malignant cell classification (Fig. 2E).

The proportions of differentiation states and meta-programs varied across organ sites (Fig. 2F). Brain and liver metastases were predominantly melanocytic (77% and 90%), whereas lung metastases carried the largest dedifferentiated fraction (neural-crest-like 28%) and, in addition, a rare neural crest-like/mesenchymal state (n = 655; 1% of malignant nuclei) that was almost exclusively found in the lung metastases (Fig. 2F). This organ pattern is, however, likely confounded by melanoma subtypes and by individual patients (Supplementary Fig. 1). The neural crest-like/mesenchymal population in particular is not a generalisable organ feature: 98% of these nuclei come from a single desmoplastic-melanoma patient (P235) and sampled on-treatment. The vast majority of cells assigned as neural crest-like came from a single patient with desmoplastic melanoma (LMM/DM). This patient’s cells are distributed across three lung samples, two pre-treatment and one on-treatment (MPI002_08, MPI002_11 and MPI002_09 respectively). This aligns with established clinical observations that desmoplastic melanomas typically lack melanocytic markers and overexpress neural crest markers [22]. Lung metastasis has been previously observed for this subtype of melanoma [23]. Our data demonstrate that the characteristics commonly observed in this subtype of primary tumour are retained in these lung metastases.

By contrast, the transitory state was distributed across all 20 patients and several subtypes and remained substantially higher in lung than in brain even when subtype was held constant (superficial spreading melanoma: 37% versus 6% transitory; Supplementary Fig. 1), pointing to a possible genuine organ contribution to dedifferentiation along the lineage axis. The melanocytic predominance of liver and brain was robust to subtype.

### Immune cell characterisation across organ sites

The 56,574 nuclei classified as Immune were used for further characterisation. After immune-specific quality control and resolution into lymphoid and myeloid identities, 45,652 lymphoid and 10,922 myeloid nuclei were retained for the sub-compartment analyses (Fig. 3A,C).

**Figure 3.**
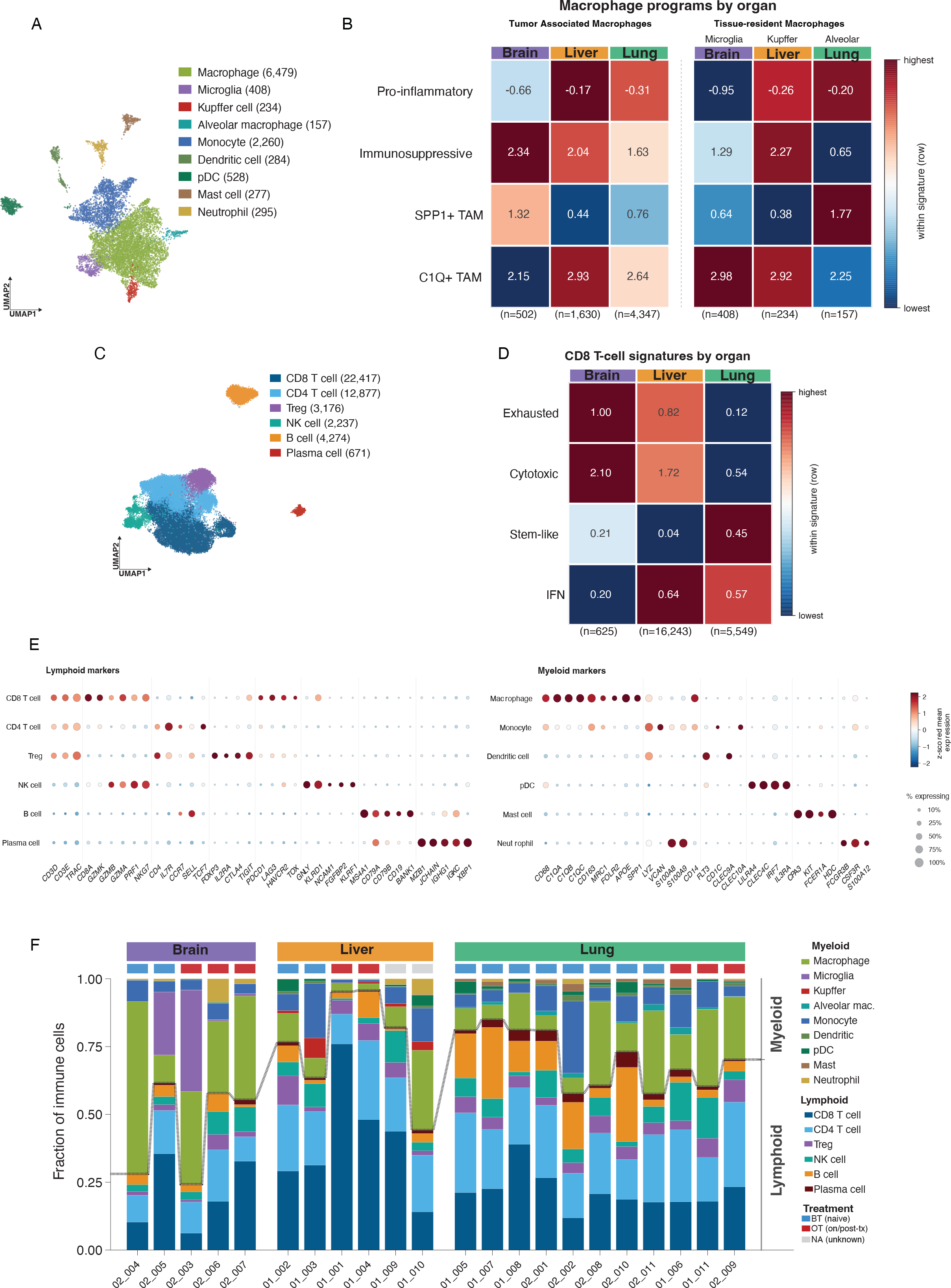
Immune cell characterisation across organ sites. **(A)** UMAP of myeloid nuclei coloured by annotated population: macrophage (6,479), microglia (408), Kupffer cell (234), alveolar macrophage (157), monocyte (2,260), dendritic cell (284), pDC (528), mast cell (277) and neutrophil (295). **(B)** Heatmaps of macrophage-program signature scores (scaled within each row) by organ. Left, tumour-associated macrophages (TAMs) across brain, liver and lung; right, tissue-resident macrophages (microglia in brain, Kupffer cells in liver, alveolar macrophages in lung). Rows: pro-inflammatory, immunosuppressive, SPP1⁺ TAM and C1Q⁺ TAM programs; sample sizes are given below each column. Brain-infiltrating myeloid cells were the most immunosuppressive and least pro-inflammatory. **(C)** UMAP of lymphoid nuclei coloured by annotated population: CD8⁺ T cell (22,417), CD4⁺ T cell (12,877), regulatory T cell (Treg; 3,176), NK cell (2,237), B cell (4,274) and plasma cell (671). **(D)** Heatmap of CD8⁺ T-cell program signature scores (scaled within each row) by organ (brain, liver, lung); rows: exhausted, cytotoxic, stem-like and IFN programs; sample sizes below each column. Brain CD8⁺ T cells scored highest for cytotoxicity, liver cells for exhaustion and IFN activity, and lung cells for the stem-like and IFN programs. **(E)** Dot plots of canonical marker genes confirming lymphoid (left) and myeloid (right) cell-type annotations. Dot size indicates the percentage of nuclei expressing each gene and colour indicates the z-scored mean expression within each population. **(F)** Stacked bar plots of the fraction of immune nuclei per sample, split into myeloid and lymphoid lineages and grouped by organ; treatment-status tiles (BT, blue; OT, red; NA, grey) are shown above each bar. Myeloid cells were relatively more abundant in brain; liver showed the strongest myeloid–lymphoid disparity (a dominant T-cell population) with high inter-patient variability; lung showed the most stable proportions across patients irrespective of treatment.

Within the myeloid compartment we annotated nine populations: macrophages, three tissue-resident macrophage subsets (microglia, Kupffer cells and alveolar macrophages), monocytes, dendritic cells, pDCs, mast cells and neutrophils (Fig. 3A) Scoring macrophages for pro-inflammatory versus immunosuppressive polarisation and for SPP1⁺ and C1Q⁺ tumour-associated-macrophage (TAM) states (Fig. 3B) showed that brain-infiltrating myeloid cells were the most immunosuppressive and least pro-inflammatory of the three organs. The three resident-macrophage subsets were each essentially organ-specific (microglia to brain, Kupffer cells to liver and alveolar macrophages to lung), recapitulating the resident myeloid identity of each organ.

Within the lymphoid compartment we annotated six populations: CD8⁺ T cells, CD4⁺ T cells, regulatory T cells (Treg), NK cells, B cells and plasma cells (Fig. 3C). Scoring CD8 T cells for cytotoxic, exhausted, stem-like and interferon (IFN) programs resolved the expected functional axes: a cytolytic-effector, exhausted, and stem-like and IFN activity (Fig. 3D).

Summarising these CD8 T-cell programs by organ revealed organ-of-residence differences in function (Fig. 3D). Brain-infiltrating cells scored highest for cytotoxicity, liver cells for exhaustion and IFN activity, and lung cells both for IFN activity and the stem-like program, suggesting that immune functional states may be shaped by the organ niche rather than being uniform across sites, although this will require validation in larger cohorts.

As a validation of our annotation, we confirmed the expression of defining markers for each immune cell type (Fig. 3E). The proportions of immune cells across patients differed widely. Generally, myeloid cells were more abundant in the brain compared to the liver and lung. The liver exhibited the most pronounced disparity between myeloid and lymphoid lineages, driven by a dominant T-cell population; however, inter-patient variability among liver samples was also substantial. Conversely, the lung constituted the most stable relative proportions of immune cells across patients regardless of treatment status.

### Stromal microenvironment characterisation across organ sites

The 18,676 nuclei classified as Stromal (grouping all non-malignant, non-immune lineages; Fig. 1), decomposing each organ’s stromal compartment into shared (pan-organ), and organ specific parenchymal populations (Fig. 4A). Pan-organ cells included fibroblasts, endothelial cells and pericytes. Liver cells were classified as hepatocytes, hepatic stellate cells and cholangiocytes; alveolar type II, alveolar type I and ciliated cells in lung; and oligodendrocytes, OPCs and neurons in brain (Fig. 4B).

**Figure 4.**
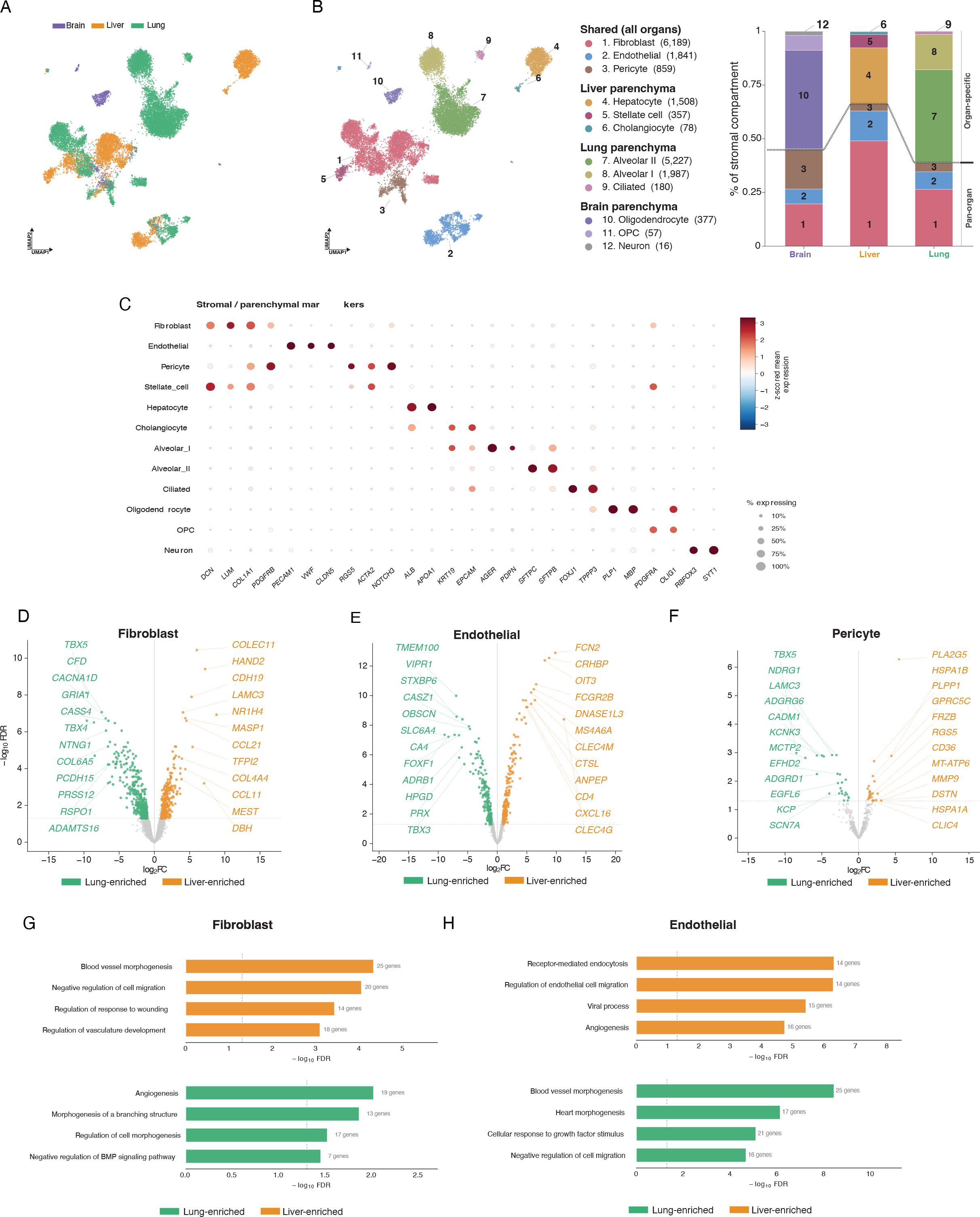
Stromal microenvironment characterisation across organ sites. **(A)** UMAP of stromal/parenchymal nuclei coloured by organ of origin (brain, liver, lung). **(B)** Left, UMAP of the same nuclei coloured by annotated population, grouped into shared pan-organ lineages: fibroblast (6,189), endothelial (1,841) and pericyte (859). Organ-specific parenchyma: liver (hepatocyte 1,508; stellate cell 357; cholangiocyte 78), lung (alveolar I 1,987; alveolar II 5,227; ciliated 180) and brain (oligodendrocyte 377; OPC 57; neuron 16). Right, stacked bar plots of the percentage of the stromal compartment attributable to pan-organ versus organ-specific populations, by organ. Brain and lung stroma were dominated by organ-specific parenchyma, whereas liver stroma was mostly the shared fibroblast/endothelial/pericyte core. **(C)** Dot plot of marker genes for each stromal and parenchymal population. Dot size indicates the percentage of nuclei expressing each gene and colour indicates the z-scored mean expression. **(D–F)** Volcano plots of differential expression between liver-enriched (orange) and lung-enriched (green) genes for each shared pan-organ population: **(D)** fibroblast, **(E)** endothelial and **(F)** pericyte. The x-axis shows log2 fold change and the y-axis −log10 FDR; selected organ-enriched genes are labelled. Lung-resident fibroblasts and pericytes expressed the lung-mesenchymal T-box factors *TBX4/TBX5*, while liver endothelium expressed a canonical liver sinusoidal endothelial cell signature (*OIT3, CLEC4G, CLEC4M, FCGR2B, DNASE1L3*). **(G, H)** Enriched GO-BP terms for liver-enriched (orange, top) and lung-enriched (green, bottom) gene sets in **(G)** fibroblasts and **(H)** endothelial cells, plotted as −log10 FDR with the number of contributing genes annotated per bar. Liver-enriched programs mapped to receptor-mediated endocytosis, endothelial-cell migration, vasculature development and wound response, whereas lung-enriched programs mapped to angiogenesis, blood-vessel and branching morphogenesis and growth-factor response. Abbreviations: BT, treatment-naïve biopsy; OT, on-/post-treatment; NA, unknown treatment status; TME, tumour microenvironment; CNV, copy-number variant; MLG, minimal-lineage gene; TAM, tumour-associated macrophage; IEG, immediate-early gene; NCSC, neural-crest stem cell; GO-BP, Gene Ontology Biological Process; FDR, false discovery rate; LSEC, liver sinusoidal endothelial cell.

Proportions using both classifications showed that brain and lung stroma were dominated by organ-specific parenchyma (glial/neural cells and alveolar epithelium, respectively), whereas liver stroma was mostly the shared fibroblast/endothelial/pericyte core (Fig 4B). Marker genes of each population are shown in (Fig. 4C).

To test whether the shared stromal lineages are themselves reprogrammed by the organ niche, we performed differential-expression analysis between organ sites for the three pan-organ populations (Fig. 4D).

Each stromal cell type carried a clear organ-specific transcriptional program (Fig. 4d– f). In fibroblasts, endothelial cells and pericytes, differential expression separates liver-enriched from lung-enriched genes. Lung-resident fibroblasts and pericytes expressed the lung-mesenchymal T-box factors *TBX4/TBX5*, while their liver counterparts up-regulated hepatic extracellular-matrix, complement and chemokine genes such as *COLEC11*, *MASP1*, *COL4A4* and *CCL21*. The contrast was sharpest in endothelium. Liver endothelial cells expressed a canonical liver sinusoidal endothelial cell (LSEC) scavenger and lectin signature (*OIT3*, *CLEC4G*, *CLEC4M*, *FCGR2B*, *DNASE1L3*), and lung endothelial cells expressed pulmonary capillary genes such as *CA4* and *FOXF1*. Together these patterns suggest that stroma at metastatic sites retains a strong imprint of its organ of residence.

Gene-ontology enrichment supported these organ-distinct programs (Fig. 4g, h). In endothelial cells, liver-enriched genes were enriched for receptor-mediated endocytosis and endothelial-cell migration, consistent with LSEC scavenger function, while lung-enriched genes were enriched for blood-vessel morphogenesis and growth-factor response. In fibroblasts, liver-enriched genes mapped to blood-vessel morphogenesis, vasculature development and wound-response terms, whereas lung-enriched genes mapped to angiogenesis, branching morphogenesis and negative regulation of BMP signalling. Because these are correlative comparisons between resident stromal populations, we describe them as organ-of-residence signatures. Whether the local microenvironment actively induces them will require functional validation studies.

## Discussion

Distant metastases remain one of the most unknown spaces of melanoma biology, where therapeutic resistance is most consequential. Existing single-cell and single-nuclei atlases of melanoma have been built almost exclusively from primary, cutaneous and lymph nodes metastases, leaving visceral metastatic lesions (brain, liver and lung) largely uncharacterised at single-nucleus resolution.

Here we present a 22 sample, 20-patient FFPE atlas of melanoma metastases distributed across these three main metastatic sites and use it to interrogate whether organ context shapes the malignant landscape and the surrounding microenvironment.

In accordance with the original snPATHO-seq report [12], we confirm that single-nucleus sequencing on archival FFPE material retains the resolution needed to recover compartment identities and detailed cell types. At the same time, it provided enough resolution to distinguish transcriptional states and meta-programs in the malignant compartment.

This makes the existing pool of clinically annotated FFPE blocks a viable substrate for atlas-scale analyses of therapeutic resistance, including retrospective cohorts. Across the 22 samples, malignant nuclei resolved into a melanocytic, transitory, neural crest-like and neural crest-like/mesenchymal differentiation axis with four overlaid functional meta-programs (cycling, hypoxia, interferon-responsive and stress), recapitulating the state framework reported in prior melanoma single-cell atlases. Several limitations should be acknowledged. The cohort size is modest (22 samples, 20 patients) and although the per-organ sample sizes are unbalanced (lung n = 11, liver n = 6, brain n = 5) the power for rigorous inter-organ statistical comparisons is limited. Organ is moreover confounded with sequencing batch (brain samples derived from a single processing batch) and with melanoma subtype (the desmoplastic, neural-crest-like tumours fall entirely in lung and, in this cohort, in a single patient), so organ-attributed malignant differences should be interpreted with caution.

We did not have matched pre-/on-treatment biopsies from the same patient (except samples MPI002_08, MPI002_09 and MPI002_11), so the BT/OT distinction is between-patient and confounded by other clinical variables. snPATHO-seq, while validated, samples nuclear RNA only and is therefore biased against transcripts whose dynamics are governed by cytoplasmic processes; some short-lived inflammatory programs may be under-detected. A separate constraint of the chemistry is that ribosomal protein genes and *HLA* genes are absent from the probe set (see Methods), which should be considered when interpreting signature scores that conventionally include these genes. Malignant-cell calls were derived from HoneyBADGER plus a GMM threshold and were not corroborated with an orthogonal CNV method (inferCNV, CopyKAT) or with spatial validation, and we did not link transcriptional findings to outcome data. The organ-of-origin imprint we describe is correlative; whether it is causal of differential drug response cannot be tested with this cohort.

In summary, this work further highlights the value of the snPATHO-seq protocol and establishes the first FFPE-compatible single-nucleus atlas of melanoma metastases across brain, liver, and lung, with harmonized cell-type, state, and meta-program annotations across organs. It will serve as a foundation for a more complete atlas and for extensive comparisons of organ-specific metastatic melanoma ecosystems.

## Methods

### Cohort and samples

Twenty-two FFPE metastatic melanoma samples from 20 patients were profiled across three organs: 5 brain, 6 liver and 11 lung samples (patient 235 contributed 3 samples). By treatment status, 12 samples were treatment-naive, 8 were on-or post treatment, and 2 were of unknown status. Written informed consent was obtained from all patients. All study procedures were in accordance with the principles of the Declaration of Helsinki, applicable Belgian law and regulations, and approved by the UZ Leuven Medical Ethical Committee (UZ Leuven protocol S67149; UAntwerpen CME-001). All available patient information is summarized in Table S1.

### Single-nucleus dissociation and sequencing

Up to two 25-µm sections per block were deparaffinized in xylene (3 × 10 min; first wash optionally at 50–55 °C), rehydrated through a graded ethanol series (100%, 70%, 50%, 30%) and washed in RPMI-1640. Tissue was digested in 1 mL RPMI-1640 with 1 mg/mL Liberase TH (45–60 min, 37 °C, 800 rpm). Nuclei were released by adding Nuclei EZ lysis buffer, pelleting (850 × g, 5 min, 4 °C) and homogenizing with a pestle in EZ lysis buffer supplemented with 2% BSA and 1 U/µL RNase inhibitor, followed by incubation on ice and, when feasible, gentle passage through a 25 G needle. Debris was removed with a 70 µm strainer; nuclei were then washed in EZ lysis buffer (2% BSA, RNase inhibitor) and twice in 0.5× PBS with 0.02% BSA, and filtered through a 40 µm strainer. Nuclei were counted by AO/PI staining on a Luna-FX7 counter, stained with DAPI, and DAPI-positive nuclei were purified by fluorescence-activated sorting on a BD FACSDiscover S8 sorter. Sorted nuclei were used immediately or cryopreserved at −80 °C in Enhancer solution (10x Genomics) with 10% glycerol. Gene expression libraries were prepared from isolated nuclei using the Chromium Fixed RNA Profiling (Flex) assay (10x Genomics) on the Chromium X. Reads were aligned to the GRCh38 reference with the snPATHO probe set and quantified with CellRanger, producing per-sample raw and filtered feature matrices. Ribosomal protein genes and *HLA* genes are not represented in the Flex probe set and are therefore not measurable by design. Antigen presentation can only be inferred from the remaining antigen-processing and loading machinery (*B2M, CD74, TAP1, TAP2, CIITA, NLRC5*).

### Initial data processing

Samples were generated in two batches (snPATHO-MPI001, n = 11; snPATHO-MPI002, n = 11). AnnData objects were assembled in Python from the per-sample CellRanger filtered matrices using Scanpy (v1.11.5).

### Quality control

Cells were filtered on three quality-control metrics: total UMI counts, number of genes detected, and percentage of mitochondrial reads (11 MT-prefixed genes). Per-sample medians of mitochondrial read percentage ranged from 0.29% to 2.57%. Cohort-wide thresholds were applied: cells with ≥ 5% mitochondrial reads were removed; cells with fewer than 200 genes detected were removed; cells above the 99.5th percentile of genes detected were removed as well as genes detected in fewer than 3 cells were removed. After quality control, the cohort was reduced from 163,616 to 154,102 cells x 18,079 genes.

### Ambient RNA assessment

Ambient RNA removal with SoupX [24] was evaluated by per-sample estimation from the raw and filtered CellRanger matrices with Leiden clustering at resolution 0.5 (estimated contamination fractions 1–9%, median 3%). Because the downstream cell-type structure was negligibly different with and without correction, while the correction altered nuclei counts and would violate the raw-count assumptions of scVI and HoneyBADGER (copy-number inference), SoupX was finally not kept in the analysis. Ambient contamination was instead handled without modifying the count matrix in two ways: ambient-dominated nuclei were flagged and removed at the annotation stage (for example, hepatocyte-ambient nuclei routed to the malignant compartment, described below), and a curated blocklist of ambient genes (*ALB, HP, FGA/B/G, APOA1/2, APOC3, ORM1, APOB, SERPINC1, TF, AHSG*) was excluded from organ-comparison differential expression.

### Doublet detection

Doublets were called per sample with three orthogonal tools: Scrublet (simulation-based), scDblFinder (gradient-boosted) and Solo (variational autoencoder, scvi-tools). A cell was flagged as a consensus doublet when at least 2 of the 3 tools agreed. Per-tool rates were 4.0% (Scrublet), 12.0% (scDblFinder) and 19.8% (Solo); the consensus doublet rate was 10.2% (15,659 cells). After removing consensus doublets, the cohort contained 138,443 cells × 18,079 genes.

### Normalisation, feature selection, dimensionality reduction

Raw counts were preserved in a dedicated layer. Counts were normalised with PFlog1pPF, with the log1p pseudocount set from a globally fitted negative-binomial overdispersion[27]. The 3,000 most highly variable genes were selected using batch-aware selection with sample identity as the batch key. Principal component analysis was computed on the highly variable gene matrix. Because PFlog1pPF is self-centering, with its final proportional-fitting step performing per-cell centering, no separate z-scaling was applied. The normalised, unscaled matrix was retained for marker detection, differential expression and visualisation.

### Data integration

Integration was performed with scVI (scvi-tools v1.4.2) on the raw counts of the 3,000 highly variable genes, using patient identity (*patient_id*) as the batch covariate and 30 latent dimensions. Training used up to 400 epochs with early stopping. A post-integration k-nearest-neighbour graph (k = 15) and UMAP were computed on the scVI latent space, and Leiden clustering was run at resolutions 0.3, 0.5, 0.8 and 1.0. Resolution 0.5 was used as the working partition for compartment classification, and all downstream visualisations use the scVI UMAP.

### Compartment classification (HoneyBADGER + signature-based GMM fusion)

Compartments (malignant, immune, stromal or uncertain) were assigned by combining gene-set scores with HoneyBADGER-derived copy-number signal in a Gaussian-mixture framework. For each cell we scored melanoma-identity signatures (Malignant_pozniak, a minimal-lineage gene signature (MLG) [6] and SuperMel_Tirosh [9]), a broad immune signature, and an organ-specific stromal/parenchymal composite (the maximum across organ-relevant stromal and parenchymal reference signatures). Parenchymal populations comprised hepatocytes, cholangiocytes and hepatic stellate cells (liver); alveolar type I, alveolar type II and ciliated cells (lung); and oligodendrocytes, OPCs and neurons (brain), alongside the shared fibroblast, endothelial and pericyte references. Gene lists were curated from CellMarker 2.0 [25] and the literature, and scoring used the Scanpy score_genes function (n_bins = 25, random_state = 42).

A high-confidence reference cell pool was built from cells scoring positively for known immune/stromal signatures and negatively for melanoma signatures. HoneyBADGER was then run against this reference to estimate per-cell mean copy-number signal (mean_cnv) and per-chromosome CNV calls. Per-feature Gaussian Mixture Models were fitted independently for maximum melanoma signature score, immune signature score, organ-specific stromal composite, mean_cnv, and per-chromosome CNV. The per-feature posteriors P(malignant | feature) were combined with a Naive Bayes log-odds rule into a single posterior P(malignant). A conservative classification was applied, cells were classified as malignant if P(malignant) > 0.90, stromal/immune (per signatures) if P(malignant) < 0.10 and uncertain otherwise. Uncertain nuclei (those with intermediate posteriors) were not force-classified but were resolved during the per-compartment annotation, and the malignant boundary was further refined by the marker-and CNV-based cleanup described below.

### Malignant cell selection and re-integration

Cells classified as malignant were subset and re-integrated with Harmony. Residual non-malignant and contaminating nuclei were removed in post-hoc cleanup using marker-gene enrichment and immune-score thresholds. Leiden clustering was re-run at resolutions 0.3, 0.5, 0.8 and 1.0, and the partition used for malignant-state analysis was chosen on stability of the clusters. Residual cross-compartment contamination was identified during malignant sub-clustering and excluded. These were nuclei that co-expressed neutrophil and monocyte markers such as *S100A8*, *S100A9*, *CSF3R* and *TREM1* alongside elevated doublet scores, while still carrying the melanocytic markers that had allowed them to pass the classifier and upstream doublet calling. Their removal yielded the 47,305 malignant nuclei of the clean atlas (Fig. 1F). Even after this removal, five nuclei belonging to a residual contamination cluster identified during malignant sub-clustering were excluded from state assignment (Fig. 2).

### Stromal / immune cell-compartment annotation (hierarchical, per sub-compartment)

The remaining nuclei were annotated through a hierarchical, iterative scVI-based pipeline. Cells were partitioned into immune, stromal/parenchymal, and uncertain sub-compartments. Each was independently re-integrated with a fresh scVI model and clustered at multiple Leiden resolutions. Cluster identity was assigned using three converging lines of evidence: Wilcoxon marker-gene rank tests; scoring against the cross-tissue snRNA-seq atlas of Eraslan et al. [26] (ArrayExpress accession E-ANND-2; top-100 author marker genes per cell type); and over-representation analysis using curated literature markers, E-ANND-2 markers and CellMarker 2.0. Each cluster’s automated suggestion was then manually reviewed against canonical lineage markers from the literature and confirmed or corrected; this curation was the final step for cell-type assignment. Resolution was chosen per sub-compartment to maximise biological interpretability.

During final atlas assembly, 6,389 of the 138,443 nuclei were flagged as low-confidence and excluded from the clean atlas. The resulting clean atlas comprises 132,054 nuclei: 47,305 malignant, 56,574 immune, 18,676 stromal/parenchymal and 9,499 uncertain. Downstream analyses used the 122,555 nuclei with a confident compartment assignment; the 9,499 uncertain nuclei are displayed in Fig. 1 but are not analysed.

### Software and reproducibility

Python analyses used scanpy (1.11.5) for normalisation, dimensionality reduction, clustering, and visualisation; scvi-tools (1.4.2) for scVI integration; scrublet for doublet detection; and decoupler (2.1.4) for pathway-set fetching. R steps (SoupX, scDblFinder, HoneyBADGER) used standard Bioconductor packages. SOLO doublet calling was run via scvi-tools.

### Data and code availability

Data and scripts are currently available upon request.

## Acknowledgments

J.C. Marine received funding from the VIB Grand Challenges Program (POINTILLISM and POINTILLISM 2.0), FWO (G052126N and G045824N), FWO-SBO (S006825N), Stichting Tegen Kanker (2022-178 and 2024-198) and iBOF (#23/005). J.P. received Stichting Tegen Kanker fellowship (2023/2310). M.P. was supported by FWO PhD fellowship (1SH7024N). P.G.D. was supported by FWO PhD fellowship (11PI824N). We acknowledge the use of the AI language model Claude (Anthropic) for assistance in troubleshooting and optimizing data analysis scripts used in the bioinformatic pipeline. We acknowledge the use of AI language model Claude (Anthropic) in the initial drafting of the text. All AI-generated text and code was rigorously reviewed, fact-checked, and substantially edited by the human authors, who assume full responsibility for the accuracy and integrity of the final manuscript. The schematic figures were generated using BioRender.com.

## Author contributions

J.C.M. and J.P. conceived the study. M.P. collected the samples and optimized and performed the single-nuclei extraction protocol. C.A.P.V. conducted the bioinformatic analyses. P.G.D. built and provided access to the biobank and offered insights regarding patient samples. F.M.B. facilitated access to samples and provided pathological expertise in selecting regions for sequencing. P.V. facilitated access to samples. O.B. Clinical expertise. C.A.P.V., M.P., J.C.M. and J.P. wrote the manuscript.

