## Supplementary Figure 1 for "Mapping tumor microenvironment heterogeneity of human melanoma metastases across distant organs"

Supplementary Fig. 1 Malignant state and meta-program composition by melanoma subtype

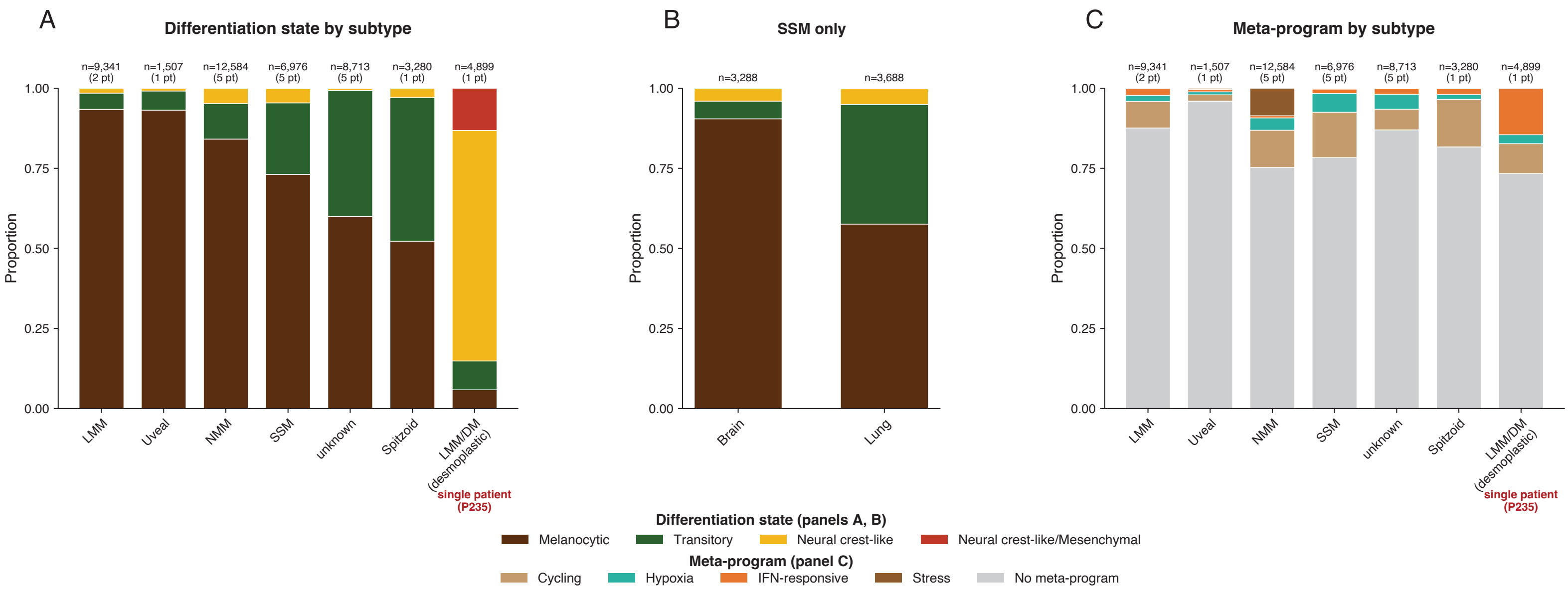

**Supplementary Figure 1. Malignant differentiation-state and meta-program composition is partially confounded by melanoma subtype and by individual patients.**

Stacked bar charts show the proportion of malignant cells assigned to each differentiation state or meta-program, grouped by melanoma histological subtype. In all panels bars are 100% stacked proportions and numbers above each bar give the number of malignant cells.

**(A)** Differentiation-state composition (melanocytic, transitory, neural crest-like, neural crest-like/mesenchymal) across melanoma subtypes. The neural crest-like state is almost entirely restricted to the desmoplastic subtype (LMM/DM), which in this cohort is contributed by a single patient (P235): 74% of all neural-crest-like cells and 98.5% of neural crest-like/mesenchymal cells derive from this patient, whose tumour cells span three lung samples (MPI002_08 and MPI002_11, pre-treatment; MPI002_09, on-treatment).

**(B)** Restricting to the superficial spreading melanoma (SSM) subtype, the transitory state remains substantially higher in lung than in brain metastases (37% versus 6%).

**(C)** Meta-program composition (Cycling, Hypoxia, IFN-responsive, Stress, and cells with no assigned meta-program) across the same subtypes.

**Abbreviations:** *LMM, lentigo maligna melanoma; LMM/DM, desmoplastic (lentigo maligna) melanoma; NMM, nodular malignant melanoma; SSM, superficial spreading melanoma.*
